# Evaluation of Reproducible and Transparent Research Practices in Sports Medicine Research: A Cross-sectional study

**DOI:** 10.1101/773473

**Authors:** Sheridan Evans, Ian A. Fladie, J. Michael Anderson, Daniel Tritz, Matt Vassar

## Abstract

**Background:** In recent years, urgency has been placed on the “reproducibility crisis” facing biomedical research. Despite efforts toward improvement, certain elements needed to reproduce a study are often lacking from publications. The current state of reproducibility within the sports medicine research community remains unknown.

**Purpose:** Our study sought to evaluate the presence of eight indicators of reproducibility and transparency to determine the current state of research reporting in sports medicine research.

**Study Design:** Cross-sectional review

**Methods:** Using the National Library of Medicine catalog, we identified 41 MEDLINE-indexed, English language sports medicine journals. From the 41 journals, we randomly sampled 300 publications that were recorded on PubMed as being published between January 1, 2014, and December 31, 2018. Two investigators extracted data in duplicate and blinded fashion.

**Results:** Of the 300 publications sampled, 280 were accessible and were screened for empirical data. Studies that lack empirical data were excluded from our analysis. Of the remaining 195 with empirical data, 10 (5.13%) publications provided data availability statements, 1 (0.51%) provided a protocol, 0 (0.0%) provided an analysis script, and 9 (4.62%) were pre registered.

**Conclusion:** Reproducibility and transparency indicators are lacking in sports medicine publications. The majority of publications lack the necessary resources for reproducibility such as material, data, analysis scripts, or protocol availability. While the current state of reproducibility cannot be fixed overnight, we feel combined efforts of data sharing, open access, and verifying disclosure statements can help to improve overall reporting.

## Introduction

According to a Center for Disease Control and Prevention (CDC) survey, participation in sports and fitness activities increased by 4 million individuals from 2014 to 2015 with a potential for increasing the risk of injury[1]. The CDC further estimated 8.6 million sports-related injuries per year from 2011-2014[1]. Concurrent to increases in sports related injuries, research publications in sports medicine have experienced considerable growth. One study reported that publications in the *American Journal of Sports Medicine* more than doubled from 2003-2013[2]. Sports medicine research has also shown signs of more robust study designs, having shifted from retrospective observational studies toward more prospective, randomized, controlled, and blinded trials[2,3]. With the increasing number of publications and higher levels of evidence, a need exists to ensure that studies are reproducible and transparent.

Reproducibility refers to “the ability of an investigator to replicate the results of a study using the same materials and procedures as the original investigators”[4]. Increased urgency has been placed on the “reproducibility crisis” currently facing biomedical research. Despite efforts toward improvement, certain elements needed to reproduce a study are often lacking from publications, including data sets, analysis codes, and software used for analysis[5]. Reproducibility is necessary for research findings to be believable and informative.[5] Replicated research can confirm that the prior outcomes were not simply a result of confirmation bias (analyzing results in a way that is partial to one’s already existing belief[6]), p-hacking (analyzing data in different ways until a nonsignificant results becomes significant[7]), or error. Some authors have encouraged replication studies in sports medicine to improve the translation of research to practice, such as within the context of the Applied Research Model for the Sport Sciences [8].

To improve confidence in sports medicine research and, thus, translation of research to practice, reproducible and transparent research practices are needed. The current climate for reproducibility within the sports medicine research community remains unknown. To address this gap, we evaluate sports medicine literature using specific markers of reproducibility and transparency.

## Methods

### Study design

We used a cross-sectional study design to evaluate specific indicators of reproducibility and transparency in sports medicine research. The study methodology is similar to that of Hardwick *et al.*[9] with minor adjustments. When applicable, the Preferred Reporting for Systematic Reviews and Meta-Analyses (PRISMA) guidelines were utilized[10]. This investigation was not subject to institutional review board oversight as no human subjects participated in our study. In order to be transparent and reproducible, information such as protocols, raw data, training recording, and additional material are available on Open Science Framework (https://osf.io/n4yh5/).

### Journal and Publication Selection

The National Library of Medicine (NLM) catalog was searched for sports medicine journals on June 5, 2019 by DT for the subject terms tag “Sports Medicine[ST]”. Inclusion criteria required journals to be MEDLINE indexed that provided full-text publications in English. The journals meeting inclusion criteria included the electronic International Standard Serial Number (ISSN) or the linking ISSN if the electronic was not identified (https://osf.io/mgczu/). PubMed was searched using the list of ISSN and filtered to include publications from January 01, 2014 through December 31, 2018. A random sample of 300 publications were included in the analysis(https://osf.io/m236a/).

### Data Extraction Training

DT lead an in-person training session for investigators assigned to extract data (SE and IF). The training reviewed study design, protocol, and data extraction. To ensure reliability of data extraction, both investigators (SE, IF) extracted data from 2 example publications and met to reconcile any discrepancies after. This same process was utilized for the first 10 sports medicine publications to further ensure reliability of extraction. After training was complete, investigators (SE, IF) extracted data from the additional 290 publications in a duplicate and blind fashion from June 11, 2019 to July 18, 2019. Investigators (SE, IF) then met to reconcile any discrepancies before extracting data from the other 290. An additional investigator was available for adjudication, but was not needed. The training session was recorded from the presenters perspective and posted online for investigator reference (https://osf.io/tf7nw/).

### Data Extraction

Our study used a Google form that was based on that used by Hardwicke *et al.* with modifications (https://osf.io/3nfa5/)[9]. We modified our form to include the 5-year impact factor and the impact factor for the most recent year listed, if available. Additional study design options were added to include cohort studies, case series, secondary analyses, chart reviews, and cross-sectional studies. Lastly, funding options were expanded to provide more insight on the specific source such as a university, hospital, public, private/industry, or non-profit. The form was then pilot-tested prior to study commencement. This form prompted investigators to look through the sample publications for information related to reproducibility and transparency. Data extracted from each publication depended on the type of study design, with studies providing no empirical data being excluded from reproducibility characteristics. Systematic reviews and meta-analyses generally do not contain the necessary data measuring materials thus excluding them from evaluating for material availability. Case reports and case series contain empirical data, but are generally not descriptive enough in their design to be reproduced in subsequent publications and were not expected to contain reproducibility characteristics[11].

### Open Access Status

We analyzed if publication’s full text was publicly available through open access during our data extraction. We used a systematic approach that first searched the Open Access Button (https://openaccessbutton.org/) using the publication title and DOI. If the website failed to find the publication publicly available or reported an error, we then searched PubMed and Google using the same identifiers as before. If the first and second step failed to find the full-text, then the publication was determined to be paywall restricted and not available through open access..

### Replication Attempts and Use in Research Synthesis

We searched Web of Science (https://webofknowledge.com) for all publications containing empirical data to determine the following: (1) the number of times a publication was cited by a systematic review/meta-analysis and (2) the number of times a publication was cited by a validity/replication study. Web of Science enabled us to sort publications by study design and to screen effectively by using the study title and abstract that was citing our original sample publication.

### Statistical analysis

We used Microsoft Excel functions to provide our statistical analysis including percentages, fractions, and confidence intervals.

## Results

### Publication selection

Our search of the NLM catalog yielded 41 eligible sports medicine journals. Our search of PubMed yielded a total of 28,921 publications from 2014 to 2018. From these publications, we extracted data from 300 randomly sampled publications. Full-text PDF versions for 280 publications (of 300) were accessible, whereas the remaining 20 publications were unavailable, and therefore excluded from our final analysis (Figure 1). Lastly, each publication was queried for the presence of indicators of reproducibility and transparency depending on the study type. Thus, Supplemental Table 1 provides further description of the eight indicators of reproducibility and transparency, their significance, and explanation of the study designs included in each analysis.

**Figure 1:**
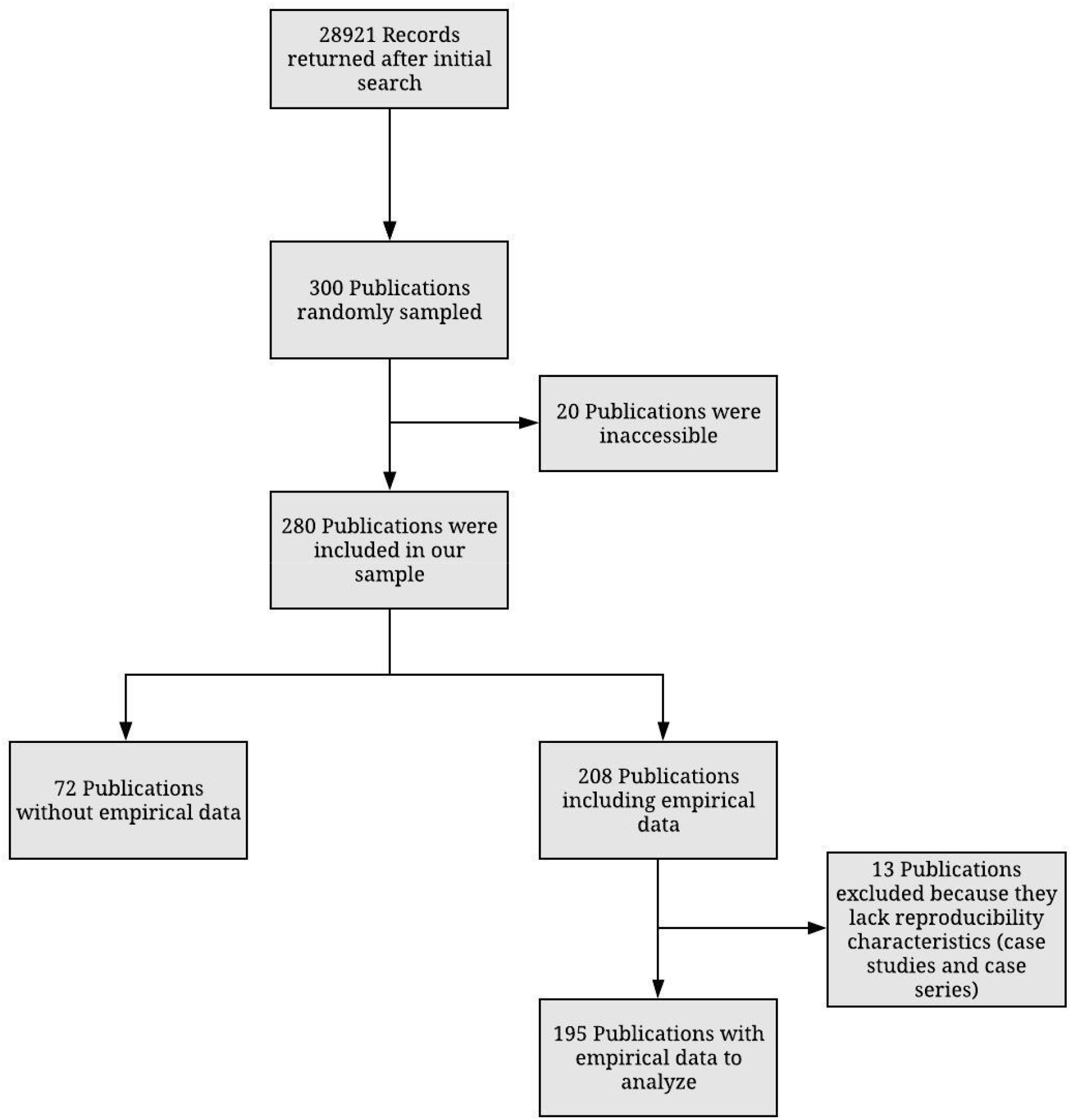
Flow diagram of Included sports medicine publications

### Sample characteristics

Clinical trials comprised the largest percentage of publications (82 of 280; 29.29%) included in our analysis. The median five-year journal impact factor was 3.279 while 31 journal impact factors were unattainable. Of the 280 accessible publications included in our random sample, 78 were publicly available using Open Access Button, Google Scholar, and/or PubMed. The remaining 202 were paywall restricted. For the purpose of our study, we considered the 20 publications that were not accessible to be restricted behind a paywall. Thus, a total of 222 publications (of 300; 74%) from our random sample were not publicly available. Additionally, almost half of the publications (129 of 280; 46.07%) had no statement regarding funding though 168 publications (of 280; 60%) provided a statement that there was no conflict of interest. Other sample characteristics can be found in Table 1.

**Table 1:**
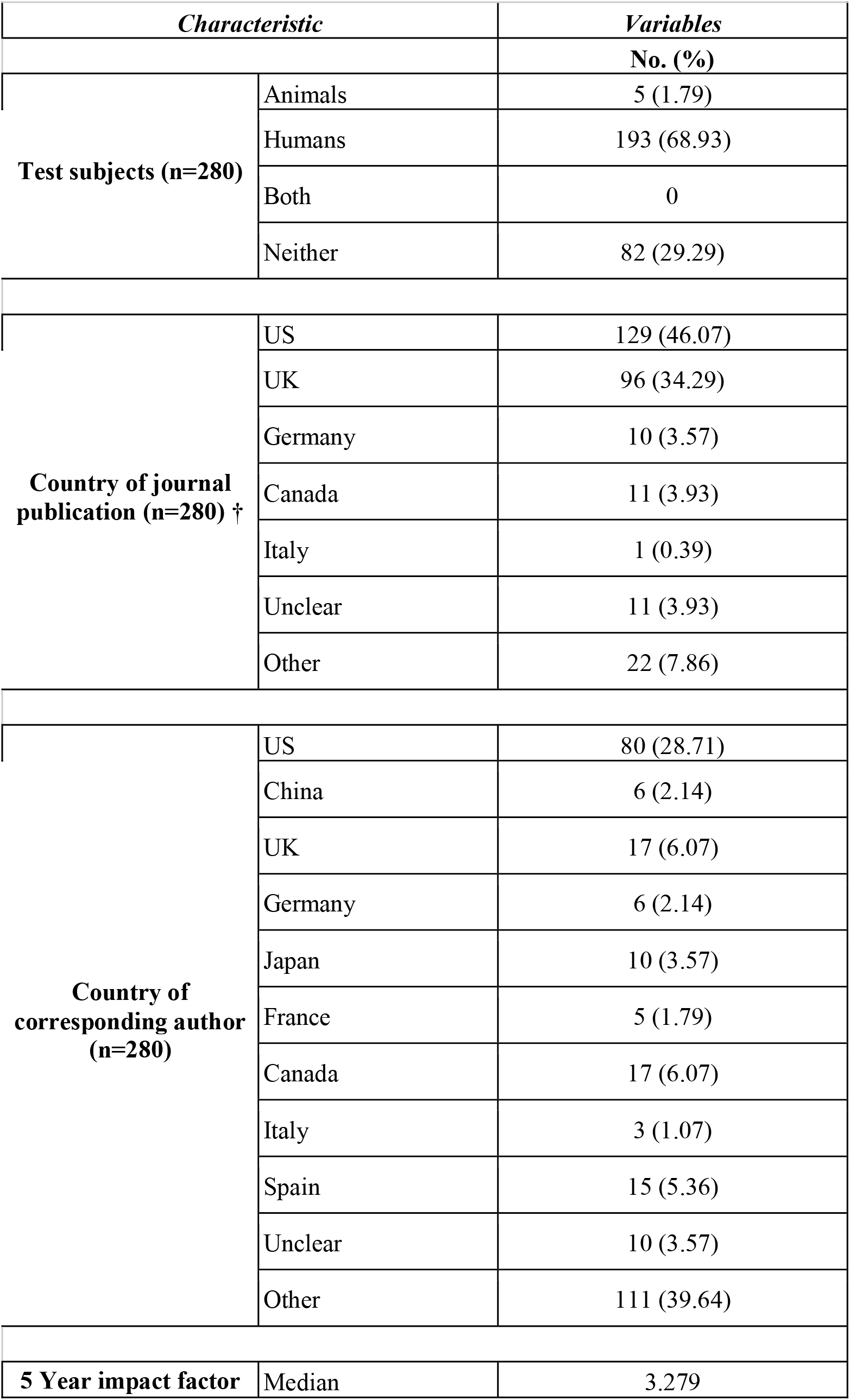
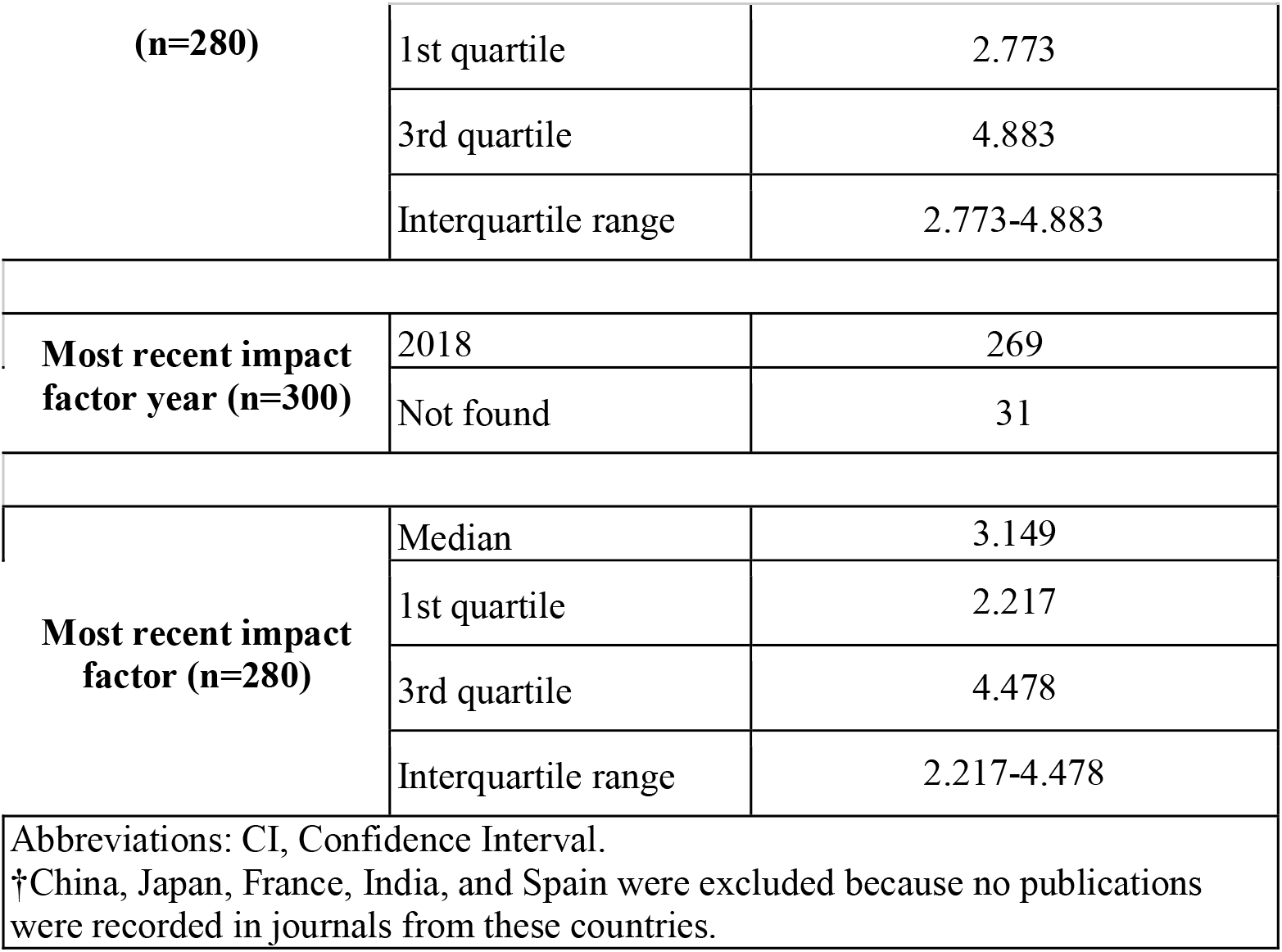
Sample characteristics of analyzed sports medicine publications

### Reproducibility components

Of 208 publications included in our analysis, only one (of 208; 0.5%) was cited in a meta-analysis or systematic review article. For further analysis we excluded 13 publications that were case studies and case series thus leaving 195 publications with empirical data. These were excluded due to their lack of reproducibility characteristics. Almost all publications lacked a data availability statement (185 of 195; 94.87%). Of the 10 (of 195; 5.13%) publications that provided a data availability statement, only one publication provided access to all raw numerical data in its unmodified state. No publication provided analysis script/code nor was any publication stated to be a replication study (0 of 195; 0%). Only one publication (of 195; 0.51%) provided a link for an accessible protocol. Our study found that 12 publications (of 195; 6.15%) included pre-registration statements, though only nine of those indicated the trial was prospectively registered. Only six of the nine pre-registrations were successfully accessed by investigators. Additionally, 8 meta-analyses were excluded from our material availability analysis. Of the remaining 187 publications, 179 (95.72%) lacked material availability statements. Additional reproducibility components can be found in Table 2 and Table 3.

**Table 2:** Reproducibility Indicators of Analyzed Orthopaedic Articles

| *Characteristics* |  | *Variables* |  |
| --- | --- | --- | --- |
|  |  | **No. (%)** | **[95% CI]** |
| **Funding (n=280)** | University | 11 (3.93) | [1.73-6.13] |
|  | Hospital | 2 (0.71) | [0.00-1.67] |
|  | Public | 49 (17.50) | [13.20-21.80] |
|  | Private/Industry | 16 (5.71) | [3.09-8.34] |
|  | Non-profit | 2 (0.71) | [0.00-1.67] |
|  | No statement listed | 129 (46.07) | [40.43-51.71] |
|  | No funding received | 41 (14.64) | [10.64-18.64] |
|  | Mixed funding received | 30 (10.71) | [7.21-14.21] |
| **Conflict of Interest statement (n=280)** | Statement provided one or more conflicts of interest | 23 (8.21) | [5.11-11.32] |
|  | Statement provided no conflict of interest | 168 (60.0) | [54.46-65.54] |
|  | No conflict of interest statement provided | 89 (31.79) | [26.52-37.05] |
| **Data availability (n=195)** | Statement was provided saying some data was available | 10 (5.13) | [2.63-7.62] |
|  | Statement was provided saying no data was available | 0 (0.0) | [0.0-0.0] |
|  | No data availability statement provided | 185 (94.87) | [92.38-97.37] |
| **Material availability (n=187)** | Statement provided saying some materials were available | 8 (4.28) | [1.99-6.57] |
|  | Statement provided saying some materials were not available | 0 (0.0) | [0.0-0.0] |
|  | No materials availability statement provided | 179 (95.72) | [93.43-98.01] |
| **Protocol availability (n=195)** | Protocol was made available | 1 (0.51) | [0.0-1.32] |
|  | No protocol was made available | 194 (99.49) | [98.68-100.0] |
| **Analysis script availability (n=195)** | Statement provided saying an analysis scripts was available | 0 (0.0) | [0.0-0.0] |
|  | Statement provided saying analysis scripts were not available | 0 (0.0) | [0.0-0.0] |
|  | No analysis script availability statement was provided | 195 (100.0) | [100.0-100.0] |
| **Replication studies (n=195)** | Replication study | 0 (0.0) | [0.0-0.0] |
|  | No statement was provided stating the study was a replicated publication | 195 (100.0) | [100.0-100.0] |
| **Cited by systematic review/Meta-analysis (n=208)** | Not cited | 162 (77.88) | [73.19-82.58] |
|  | Cited a single time | 19 (9.13) | [5.87-12.39] |
|  | Cited one to five times | 25 (12.02) | [8.33-15.70] |
|  | Cited greater than five times | 2 (0.96) | [0.00-2.07] |
| **Cited by a replication study (n=208)** | Not cited | 207 (99.52) | [98.74-100.0] |
|  | Cited a single time | 1 (0.48) | [0.0-1.26] |
| **Pre-registration (n=195)** | Statement provided saying publication was pre--registration | 9 (4.62) | [2.24-6.99] |
|  | Statement provided saying publication was not pre--registration | 3 (1.54) | [0.15-0.29] |
|  | There is no pre--registration statement provided in the publication | 183 (93.85) | [91.12-96.57] |
| **Open access (n=300)** | Yes the publication was found via Open Access Button | 74 (24.67) | [19.79-29.54] |
|  | Yes the publication was found via other means | 4 (1.33) | [0.0-2.63] |
|  | The publication could not access<br>through paywall | 202 (67.33) | [62.03-72.64] |
| Abbreviations: CI, Confidence Interval. |  |  |  |

**Table 3:**
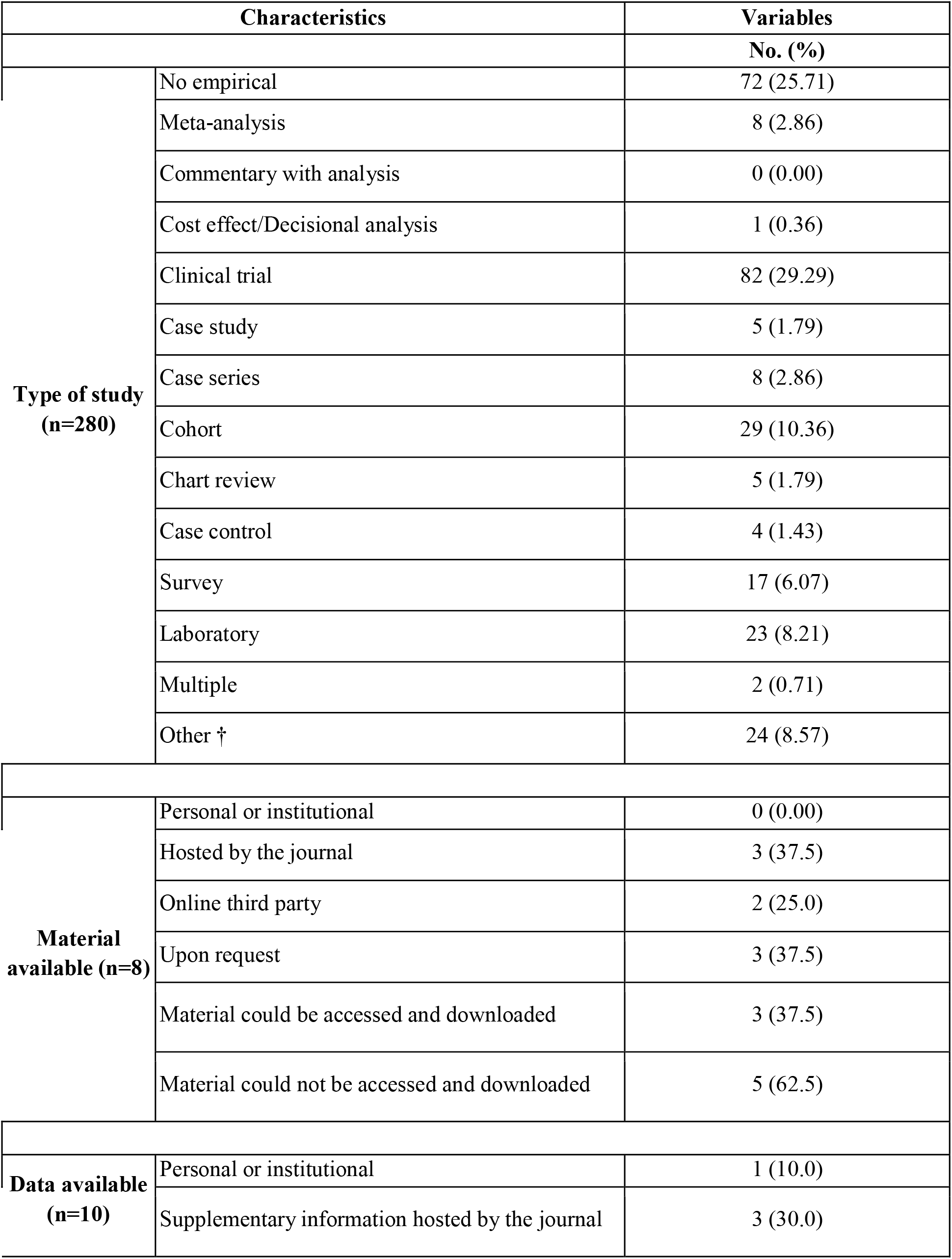
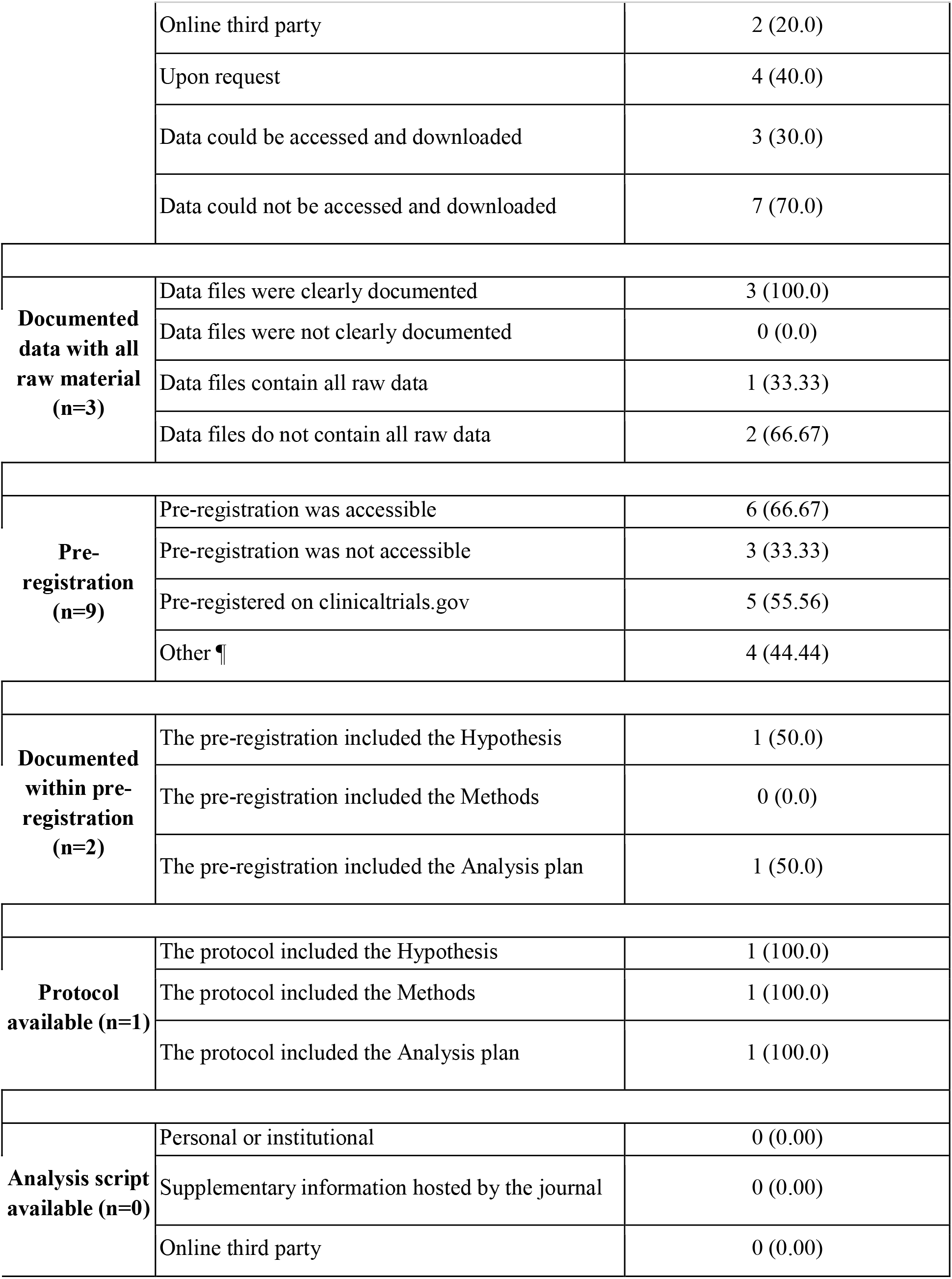
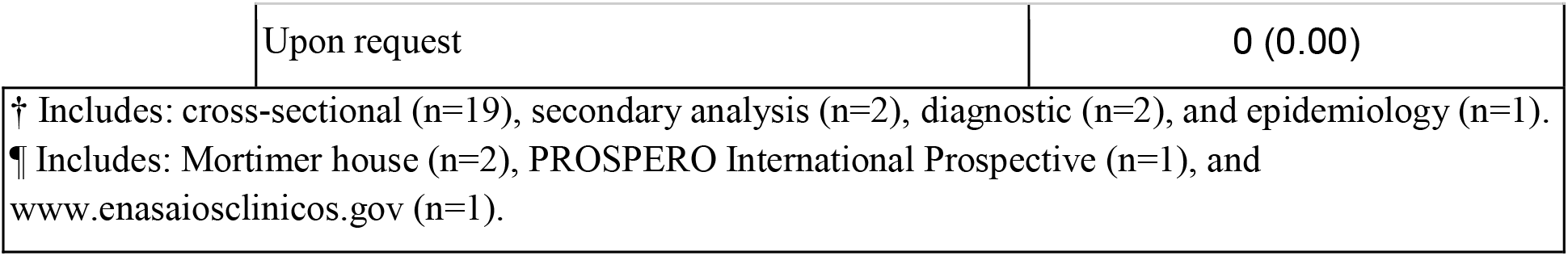
Additional Reproducibility Characteristics

## Discussion

Our study found that the majority of sports medicine publications lack the necessary resources for reproducibility. Few publications provided statements regarding material, data, analysis scripts, or protocol availability. Furthermore, less than half of our publications were available as open access. These factors are critical for reproducibility of scientific research. For example, both raw data and analysis scripts are considered to be minimum components necessary for reproducibility[5]. Hardwicke *et al.* found that of 198 social sciences studies, 8% provided raw data, 3% provided analysis scripts, and none provided protocols. In order to increase reproducibility and transparency, we discuss the importance of data sharing, open access publishing, and disclosure of financial conflicts of interests. In this discussion, we contrast the policies of 3 journals throughout — *British Journal of Sports Medicine*, *Sports Medicine*, and *American Journal of Sports Medicine* (selected from having the highest impact factors among sports medicine journals) — to showcase variation of current practices within the specialty. As journals are the final arbiters of what research is published and disseminated, we focus on them here. Clearly, we acknowledge that other stakeholders also play a role in facilitating more reproducible research.

First, data sharing is important for study reproducibility and credibility, but our findings demonstrate a lacking within sports medicine literature[12]. The International Committee of Medical Journal Editors (ICJME) recommends authors make raw data publicly available as it can contribute to the confidence in findings, ability to be validated results by independent researchers, and enable reproducibility studies to be accurately conducted[13]. However, difficulties have arisen with the complexity and varying forms of data available to authors. To combat these complications, journals should provide authors with increased resources and recommendations to improve the likelihood of making raw data publicly available[14]. The *British Journal of Sports Medicine* (*BJSM*) follows tier 2 of the *BMJ* which requires a data sharing plan be preregistered for clinical trials with raw data being available upon reasonable request and strongly recommends public access to supporting data for any study design[15,16]. The *Sports Medicine* journal does not explicitly require any raw data but states that it must be presented if requested by editors. This journal recommends the use of data repositories to make raw data publicly available and encourages including a data availability statement that describes if and where data are available[17]. Lastly, *The American Journal of Sports Medicine (AJSM)* does not specify any data sharing requirements as terms for publications and only mentions that authors may be asked to supply raw data by editors, which can lead to rejection if not provided[18]. These top journals in sports medicine have varying policies regarding data availability and should strive to encourage or require authors to make raw data publicly available when possible to increase the reproducibility and transparency of study findings.

Second, open access has numerous benefits, including more views and citation counts[19]. On the other hand, criticisms have been raised that open access publishing promotes publications from first world countries and limits publication to authors who have funding available for article processing charges, which in most cases, are substantial[20].For example, *BJSM* has an open access option, which, at the time of this writing, costs 1,950 GBP. *Sports Medicine*, published by Springer, offers Open Choice which costs 2,490 GBP, while no information is provided for the cost of open access for *AJSM*. In cases where sister open access journals have been created, it may be viewed that research studies which do not pass muster due to a lack of novelty or importance, may be suitable only for open access publication in the lower tiered journals. Another consideration with respect to open access publication is the proliferation of predatory publishers and journals. These journals often have names that closely resemble established journals and can mislead authors into submitting research manuscripts to them. While our views on open access publishing are generally positive, we recognize the inherent limitations and criticisms regarding it.

Financial conflicts of interest are consequential when study authors receive payments from companies, with these interests having the potential to bias study outcomes. Yet, FCOI declarations may not always be accurate. In one study, we found that approximately one-third of authors of pivotal oncology trials inaccurately disclosed industry payments. The ICMJE requires broad disclosure of FCOI for 36 months. Under these guidelines, authors must disclose all interests relevant to the present work as well as those outside of the present work. In sports medicine, financial disclosure policies and procedures also differed between journals. The *AJSM* requires disclosure of industry affiliations for a 5 year period and checks all disclosures against records from the Center for Medicare and Medicaide’s Open Payments Database. The *BJSM* and *Sports Medicine* requires that each author fill out the ICMJE conflict of interest form and submit it for review at the same time as the manuscript[17,21]. We recommend that other journals within sports medicine consider adoption of the Open Payments Database for verification of disclosures. At the very least, future studies should be conducted to determine the effects of using this database on the accuracy of author disclosure.

### Moving forward

The poor state of reproducibility in sports medicine research may be partially caused by lack of proper training for investigators. It is imperative that investigators have proper training in quality reporting and the components necessary for reproducibility. Courses are emerging for this type of training due to increasing reproducibility crisis. For example, as part of their movement to increase rigor and reproducibility, the NIH provides both webinars and modules to train investigators on such topics. Training includes core issues in research such as lack of transparency, analysis approaches, and review of recent practices. All training modules and webinars, while not comprehensive training, are available online and free to the public.[22] We recommend primary investigators train their teams to ensure proper techniques are used to increase reproducibility of their research.

Perhaps most importantly, research as a whole must collaborate to reduce the bias against replication studies in order to see an increase in reproducibility. Researchers may not provide all components necessary for reproducibility because replication studies are conducted less frequently. For example, a study by Makel et al. analyzed 500 randomly selected psychological research publications from 1900 to 2012 and found replication studies to be published at an overall rate of only 1.07%[23]. A possible reason for the lack of replication studies is that researchers believe there is editorial bias against such studies. In a questionnaire given to both editors and reviewers, 74% of editors and 54% of reviewers said they believed novel study results to be more important than that of replication studies[24]. Cassey et al. suggests three reasons why reproducibility and replicability is important: to protect against misrepresentation of results, data loss and human error, and deliberate fraud[25]. Replicated studies could provide safeguards against these potential errors while giving clinicians confidence in the credibility and reliability of research outcomes. The field of sports medicine is leading the way with a call for replication studies within the Applied Research Model for the Sport Sciences model. This model acknowledges the bias against replication studies along with the importance of novel research and proposes that researchers attempt to replicate previous findings within their own novel studies as a solution[8]. Thus, we recommend an increased level of collaboration in the scientific community to encourage the publication of replication studies, directly increasing reproducibility and reliability of the literature.

### Strengths and Limitations

One strength of our study was using Cochrane gold standard data extraction in a blinded and duplicate fashion to minimize bias and ensure accuracy[26]. Additionally, to our knowledge this was the first study to address reproducibility and transparency indicators in sports medicine. To encourage reproducibility, we provided all data, training, and methods on Open Science Framework. Concerning limitations, the time frame of our search and the selected journals may limit the generalizability of our findings. Additionally, our data extracted was limited to what was presented within the full-text publication and did not contact authors about the possibility of sharing information with us. Had we contacted authors and requested access to the measures of this study, the data could potentially be higher. We thought this was unlikely as authors often do not respond to requests or deny access to the requested materials[27].

### Conclusion

The poor state of reproducibility is far reaching in sports medicine research. Steps must be taken to improve reproducibility and transparency as our study found very few of the publications in sports medicine providing the necessary materials for reproducing study findings. The recommendations made in this study should foster an improved focus on reproducibility.

## Disclosures

The authors report no conflicts of interest.

## Author contributions

MV and DT conceived the study and designed the protocol. MV and DT supervised the study. MV received research funding. SE and IF extracted data. SE and IF conducted data analysis and organization. MA conducted the final data analysis. SE, IF, MA, DT, and MV drafted the manuscript. All authors provided critical feedback, ideas, and editing for the manuscript and have approved the final version. SE assumes responsibility for this manuscript.

## Funding

This study was funded by the 2019 Presidential Research Fellowship Mentor – Mentee Program at Oklahoma State University Center for Health Sciences.

---

**Supplemental table 1:**
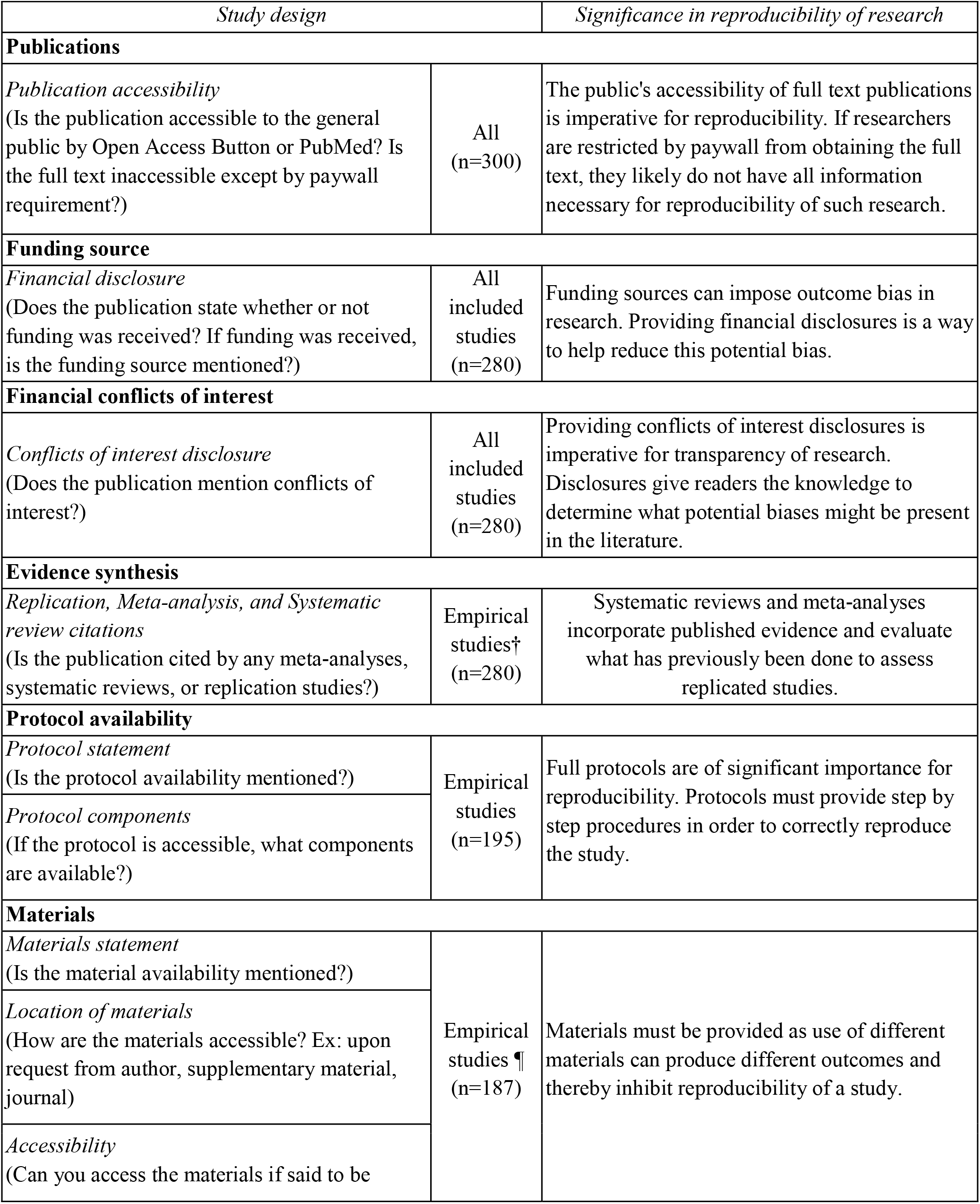
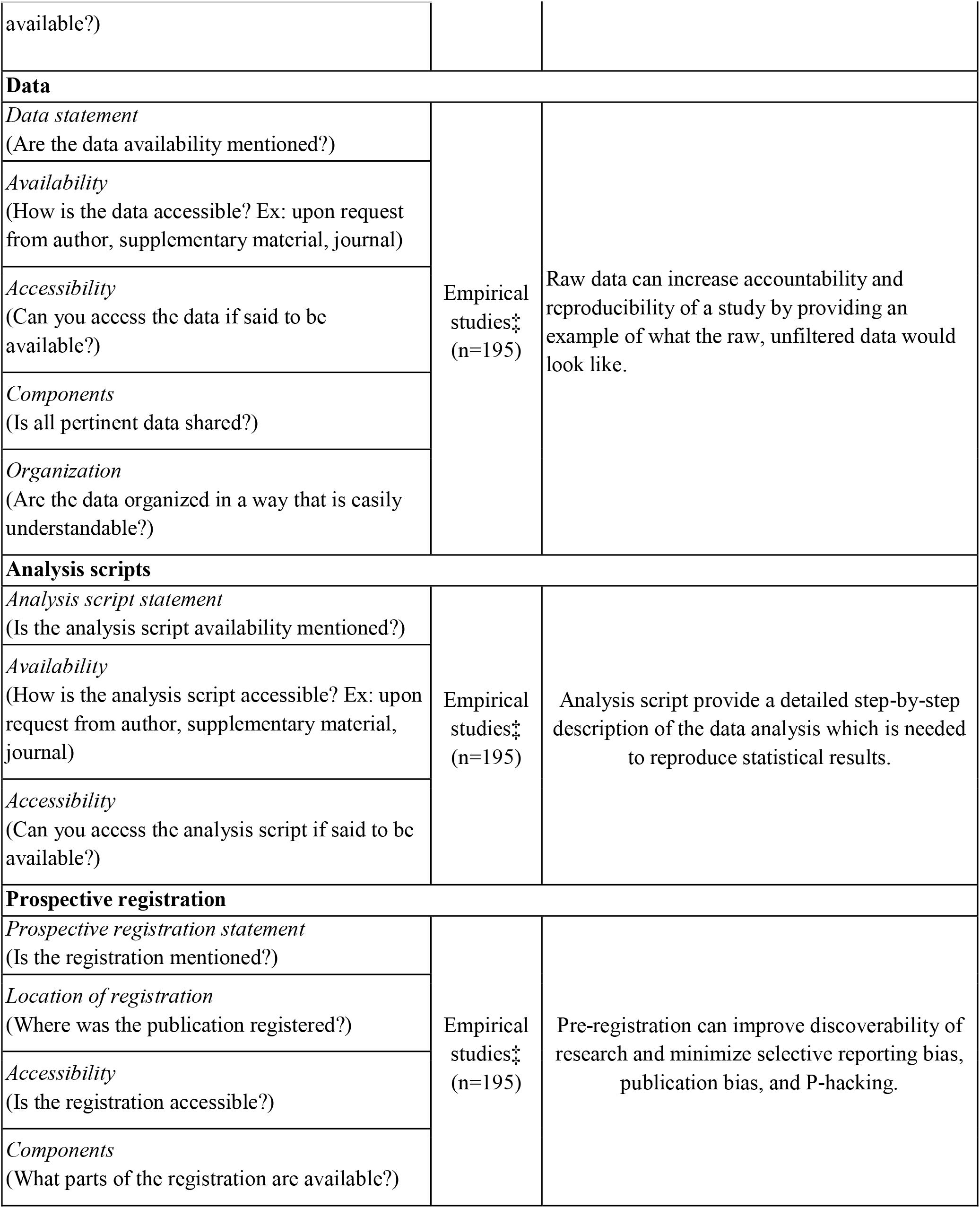
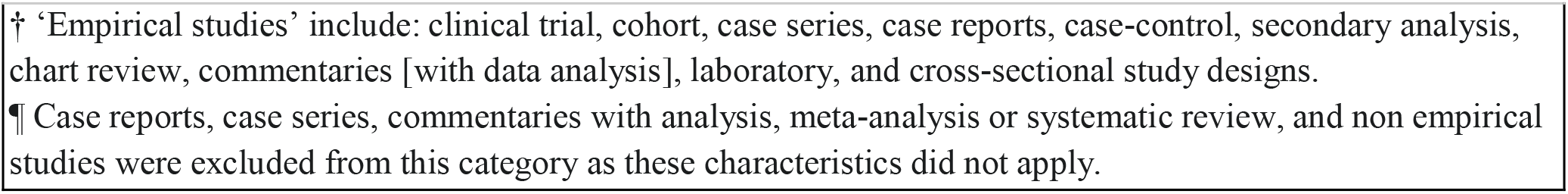
Analyzed components of each publication. Components analyzed varied by study type. Additional details can be found at: https://osf.io/x24n3/

| Study design |  | Significance in reproducibility of research |
| --- | --- | --- |
| Publications |  |  |
| Publication accessibility<br>(Is the publication accessible to the general public by Open Access Button or PubMed? Is the full text inaccessible except by paywall requirement?) | All<br>(n=300) | The public's accessibility of full text publications is imperative for reproducibility. If researchers are restricted by paywall from obtaining the full text, they likely do not have all information necessary for reproducibility of such research. |
| Funding source |  |  |
| Financial disclosure<br>(Does the publication state whether or not funding was received? If funding was received, is the funding source mentioned?) | All included studies<br>(n=280) | Funding sources can impose outcome bias in research. Providing financial disclosures is a way to help reduce this potential bias. |
| Financial conflicts of interest |  |  |
| Conflicts of interest disclosure<br>(Does the publication mention conflicts of interest?) | All included studies<br>(n=280) | Providing conflicts of interest disclosures is imperative for transparency of research. Disclosures give readers the knowledge to determine what potential biases might be present in the literature. |
| Evidence synthesis |  |  |
| Replication, Meta-analysis, and Systematic review citations<br>(Is the publication cited by any meta-analyses, systematic reviews, or replication studies?) | Empirical studies†<br>(n=280) | Systematic reviews and meta-analyses incorporate published evidence and evaluate what has previously been done to assess replicated studies. |
| Protocol availability |  |  |
| Protocol statement<br>(Is the protocol availability mentioned?) | Empirical studies<br>(n=195) | Full protocols are of significant importance for reproducibility. Protocols must provide step by step procedures in order to correctly reproduce the study. |
| Protocol components<br>(If the protocol is accessible, what components are available?) |  |  |
| Materials |  |  |
| Materials statement<br>(Is the material availability mentioned?) | Empirical studies ¶<br>(n=187) | Materials must be provided as use of different materials can produce different outcomes and thereby inhibit reproducibility of a study. |
| Location of materials<br>(How are the materials accessible? Ex: upon request from author, supplementary material, journal) |  |  |
| Accessibility<br>(Can you access the materials if said to be |  |  |
| available?) |  |  |
| **Data** |  |  |
| *Data statement*<br>(Are the data availability mentioned?) | Empirical studies‡<br>(n=195) | Raw data can increase accountability and reproducibility of a study by providing an example of what the raw, unfiltered data would look like. |
| *Availability*<br>(How is the data accessible? Ex: upon request from author, supplementary material, journal) |  |  |
| *Accessibility*<br>(Can you access the data if said to be available?) |  |  |
| *Components*<br>(Is all pertinent data shared?) |  |  |
| *Organization*<br>(Are the data organized in a way that is easily understandable?) |  |  |
| **Analysis scripts** |  |  |
| *Analysis script statement*<br>(Is the analysis script availability mentioned?) | Empirical studies‡<br>(n=195) | Analysis script provide a detailed step-by-step description of the data analysis which is needed to reproduce statistical results. |
| *Availability*<br>(How is the analysis script accessible? Ex: upon request from author, supplementary material, journal) |  |  |
| *Accessibility*<br>(Can you access the analysis script if said to be available?) |  |  |
| **Prospective registration** |  |  |
| *Prospective registration statement*<br>(Is the registration mentioned?) | Empirical studies‡<br>(n=195) | Pre-registration can improve discoverability of research and minimize selective reporting bias, publication bias, and P-hacking. |
| *Location of registration*<br>(Where was the publication registered?) |  |  |
| *Accessibility*<br>(Is the registration accessible?) |  |  |
| *Components*<br>(What parts of the registration are available?) |  |  |

† ‘Empirical studies’ include: clinical trial, cohort, case series, case reports, case-control, secondary analysis, chart review, commentaries [with data analysis], laboratory, and cross-sectional study designs.
¶ Case reports, case series, commentaries with analysis, meta-analysis or systematic review, and non empirical studies were excluded from this category as these characteristics did not apply.

